# Identification of Cancer–associated Metabolic Vulnerabilities by Modeling Multi-objective Optimality in Metabolism

**DOI:** 10.1101/198333

**Authors:** Ziwei Dai, Liyan Xu, Hongrong Hu, Kun Liao, Shuye Deng, Qiyi Chen, Shiyu Yang, Qian Wang, Shuaishi Gao, Bo Li, Luhua Lai

## Abstract

Computational modeling of the genome-wide metabolic network is essential for designing new therapeutics targeting cancer-associated metabolic disorder, which is a hallmark of human malignancies. However, previous studies generally assumed that metabolic fluxes of cancer cells are subjected to the maximization of biomass production, despite the wide existence of trade-offs among multiple metabolic objectives. To address this issue, we developed a multi-objective model of cancer metabolism with algorithms depicting approximate Pareto surfaces and incorporating multiple omics datasets. To validate this approach, we built individualized models for NCI-60 cancer cell lines, and accurately predicted cell growth rates and other biological consequences of metabolic perturbations in these cells. By analyzing the landscape of approximate Pareto surface, we identified a list of metabolic targets essential for cancer cell proliferation and the Warburg effect, and further demonstrated their close association with cancer patient survival. Finally, metabolic targets predicted to be essential for tumor progression were validated by cell-based experiments, confirming this multi-objective modelling method as a novel and effective strategy to identify cancer-associated metabolic vulnerabilities.

## Introduction

Since Otto Warburg’s first description that cancer cells exhibit abnormally high glucose uptake and concomitant lactate secretion(Warburg, 1956), metabolic alteration is widely noted as a hallmark of cancer(Boroughs and DeBerardinis, 2015; DeBerardinis and Chandel, 2016; Hanahan and Weinberg, 2011; Pavlova and Thompson, 2016). Besides the “wasteful” metabolism known as aerobic glycolysis or the Warburg effect(Dai et al., 2016; Liberti and Locasale, 2016), metabolism in malignant cells are shifted at the systematic level due to numerous factors including nutrient and oxygen availability in the tumor microenvironment, materials and energy required for rapid cell proliferation, as well as oncogenic signaling pathways. Thus, targeting metabolic reprogramming in cancer is a promising strategy for designing anti-tumor therapeutics(Cheong et al., 2012; Martinez-Outschoorn et al., 2017; Vander Heiden, 2011; Vernieri et al., 2016).

While traditional methods are suited to dissect limited numbers of metabolic pathways, systems biology is a powerful tool to study metabolism from a global perspective(Yizhak et al., 2015). Within the field of cancer metabolism, analyses of genome-scale metabolic models (GSMMs)(Thiele and Palsson, 2010; Thiele et al., 2013) enabled researchers to elucidate the plausible mechanism of Warburg effect(Shlomi et al., 2011), quantify efficacies and side effects of cancer therapeutics(Agren et al., 2014; Folger et al., 2011; Shaked et al., 2016; Yizhak et al., 2014a; Yizhak et al., 2014b), and unravel context-dependent functionality of metabolic enzymes during tumor progression(Frezza et al., 2011; Megchelenbrink et al., 2015; Rabinovich et al., 2015; Tardito et al., 2015). Among various strategies, flux balance analysis (FBA) exhibits itself as a highly effective approach to analyze GSMMs(Orth et al., 2010). FBA commonly assumes that cells organize metabolic fluxes by perusing metabolic objectives subjected to certain stoichiometric constraints and upper/ lower limits. The assumption of maximized biomass production (representing for optimal cancer cell growth) has been widely used in previous studies modeling cancer metabolism.

Despite the rapid development in modeling cancer metabolism, the fundamental assumption of most computational methods – maximization of growth rate in cancer cells – is still open to doubt. Although studies investigating the metabolic objectives of cancer cells were scarce, several studies focusing on unicellular organisms provided useful insights(Gianchandani et al., 2008; Knorr et al., 2007; Schuetz et al., 2007). Interestingly, the hypothesis of single-objective metabolic optimization was challenged even in *Escherichia coli*. Comparison of experimentally-measured metabolic fluxes and the Pareto-optimal surface defined by multiple metabolic objectives revealed that cellular metabolism may be determined by trade-off among three competing objectives: maximization of biomass yield, maximization of ATP production, and minimization of gross metabolic fluxes(Schuetz et al., 2012). Similarly, the trade-off between biomass yield and ATP production was also considered as one plausible mechanism underlying tumor-associated metabolic disorders including the Warburg effect(Pfeiffer et al., 2001).

In line with these findings, we present here the first theoretical strategy involving multi-objective optimality for modeling cancer metabolism to our best knowledge. Specifically, we developed algorithms for sampling balanced flux configurations with Pareto optimality and building individualized models based on publically-available omics data. To demonstrate our methodology, we constructed individualized models for NCI-60 cancer cell lines and predicted the impact of metabolic gene ablation on Pareto optimality, metabolism, and cell viability. With this approach, we identified a list of metabolic enzymes essential for cell proliferation and aerobic glycolysis (the Warburg effect), and further validated this list through survival analysis and cell-based experiments. These metabolic hubs will likely improve our understanding of cancer-associated metabolic disorders, and provide potential targets for novel cancer therapeutics.

## Results

### Four-objective optimization model for cancer metabolism

Metabolism is pivotal for biomass synthesis and energy production indispensable for cell viability. However, these two goals contradict with each other to some extent during cell division. For instance, when glucose is completely oxidized to carbon dioxide to generate maximal amounts of ATP, it can no longer be utilized as biomass precursors for amino acid and nucleotide biosynthesis. Therefore, we reasoned that the distribution of metabolic flux can only be comprehensively determined by multiple biological objectives (Fig 1A), including (1) maximization of biomass production, which is frequently considered as the only objective in previous FBA studies of cancer cells(Folger et al., 2011; Gatto et al., 2015; Megchelenbrink et al., 2015; Yizhak et al., 2014a), (2) maximization of ATP hydrolysis, which is considered as the objective in some FBA studies of non-malignant cells(Folger et al., 2011; Yizhak et al., 2014b), (3) minimization of total abundance of metabolic enzymes, which is an analogue of the solvent capacity constraint(Shlomi et al., 2011; Vazquez et al., 2010), and (4) minimization of total carbon uptake(Savinell and Palsson, 1992). These four objectives reflect different aspects of metabolic demand, covering both maximization of biomass yield and minimization of energy cost. Combining them with a genome-scale metabolic network Recon 1(Duarte et al., 2007), we created a multi-objective linear programming model (Fig 1B), which serves as the theoretical framework for our subsequent analysis (see Materials and Methods for details).

**Fig 1.**
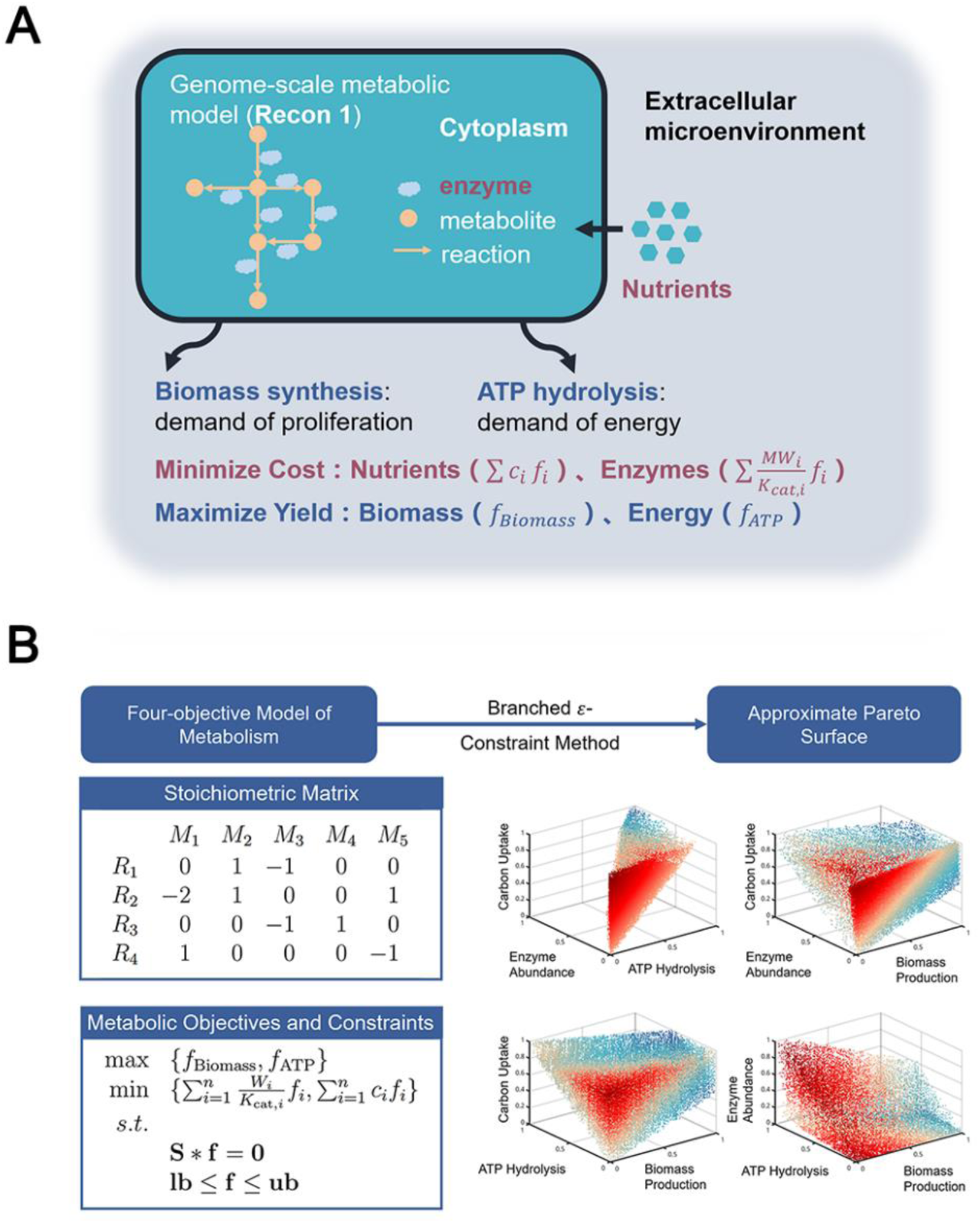
Four-objective optimization model for cancer metabolism. (A) Illustration of the four metabolic objectives incorporated in this model. (B) Mathematical description of the model and approximate Pareto surface projected on four ternary combinations of included objectives. Data points are presented in shade to depict the shape of Pareto surface.

Based on this model, we quantitatively describe the trade-off among multiple metabolic objectives by considering solutions with Pareto optimality (Fig 1B). Pareto optimality is defined by inabilities to further optimize one objective function without making any other objectives worse off. For instance, a metabolic flux configuration with Pareto optimality with regarding to the two objectives of maximizing biomass and ATP yield is one that cannot be altered to yield both higher biomass synthesis and higher ATP production. To uniformly sample from all solutions with Pareto optimality, we designed an algorithm based on ε-constraint method(Mavrotas, 2009), namely branched ε-constraint method (BECM). The resulting collection of all Pareto solutions, or the Pareto surface, can be visualized by projection to any ternary combinations of the metabolic objectives as mentioned above (Fig 1B).

### Individualized Pareto models accurately predict cell proliferation and responses to metabolic gene ablations in NCI-60 cancer cell lines

To further validate the four-objective optimization method in modeling cancer metabolism, Pareto optimality is assumed to be achieved in the examined cancer cells or tissue types, and their metabolic flux configurations could be reconstructed by searching for Pareto solutions harboring the highest consistency with protein expression and metabolic flux profiles. To do this, we developed a population-based strategy based on the assumption that for a fixed metabolic pathway, the corresponding enzymatic expression correlates with its governed metabolic flux (Fig 2A). Specifically, we used multi-omics datasets including LC-MS/MS based proteomics(Gholami et al., 2013) and consumption-release (CORE) profiles of metabolites(Jain et al., 2012) to reconstruct Pareto models for NCI-60 cell lines.

**Fig 2.**
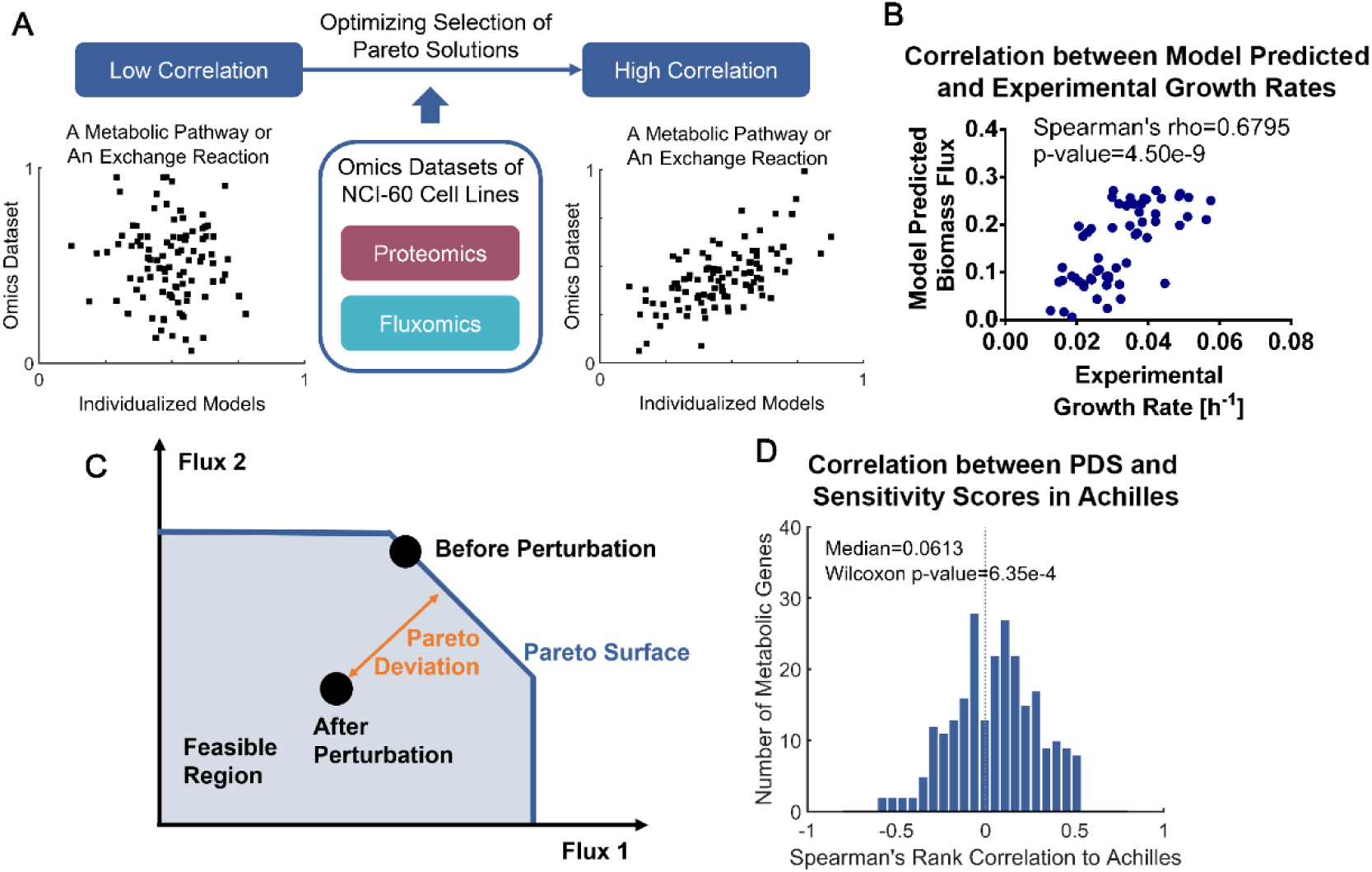
Individualized Pareto models for NCI-60 cancer cell lines predict cell proliferation rates and responses to metabolic gene ablations. (A) Illustration of the strategy used in constructing the individualized models based on multiple omics datasets. (B) Comparison between actual and model-predicted cell growth rates in the NCI-60 cancer cell panel. (C) Illustration of Pareto deviation score (PDS) as a metric quantifying the impact of metabolic perturbation on cell viability. (D) Distribution of Spearman’s rank correlation coefficients between model-predicted PDS values and sensitivity scores for metabolic gene ablations in the Achilles Database.

We then validated the Pareto models by comparing model-predicted biomass fluxes to the actual cell growth rates, and found that theoretical biomass production fluxes accurately predict the cell growth rates as previously reported (Fig 2B, Spearman’s rank correlation coefficient = 0.68). Moreover, model-predicted metabolic fluxes positively correlate with pathway-level protein expression profiles and experimentally-measured metabolic fluxes, demonstrating that our Pareto models successfully recapitulate cancer-associated metabolic phenotypes (Fig S1). Convergence to different solutions only slightly affect prediction results (Fig S2), while fewer metabolic objectives (Fig S3) or less omics data input (proteomics or CORE only, Fig S4) would significantly reduce prediction accuracies, suggesting that all four objectives and both omics datasets are indispensable for modeling the metabolic landscape of cancer cells.

Next, we applied this model to predict cellular responses to metabolic gene ablations and compared the calculated results with Achilles, a genome-scale gene essentiality database(Cowley et al., 2014). Based on the assumption of Pareto optimality in modeling cancer metabolism, the distance between a metabolic flux configuration and the Pareto surface reflects how fitness of cells bearing such flux configuration is impaired. To better quantify this deviation, we defined a Pareto deviation score (PDS) as the Euclidean distance between the flux configuration after metabolic gene knockdown and the Pareto surface (Fig 2C, Materials and Methods). For each metabolic gene registered in Achilles and associated with only one reaction in Recon 1 (245 in total), we simulated the flux configurations after its knockdown in all NCI-60 cell lines and computed the PDS values. Then we compared the resultant PDS with sensitivity scores in Achilles and demonstrated their positive correlations in a majority of examined genes (Fig 2D, p<0.001, Wilcoxon’s signed-rank test), thus corroborating the close association between theoretical deviations from Pareto optimality in achieving all four objectives and impairments of cell viability.

### Metabolic targets identified by Pareto surface analysis are essential for cancer progression

Given that model-predicted PDS values reflect sensitivities to metabolic perturbation, we next sought to identify anti-tumor metabolic targets using this novel model. It is noted that rapid proliferation and the Warburg effect (fermentation of glucose in the presence of oxygen) are essential features of cancer cells, with the latter more related to tumor metastasis and drug resistance(Gatenby and Gillies, 2004; Liberti and Locasale, 2016). Counteracting these two features are critical for developing anti-tumor therapies. Therefore, we designed a perturbation strategy leading to larger Pareto deviation in flux configurations with increased biomass production or enhanced Warburg effect, aiming to selectively impair the viability of malignant cells. This perturbation can be achieved by activation or inhibition of metabolic enzymes, which can be quantified in our model as increased or decreased metabolic fluxes governed by a particular enzyme. Without loss of generality, we were able to use cell growth rate as a representative phenotype to illustrate our strategy for target identification. Next, we projected the Pareto surface to a two-dimensional space spanned by growth rate and one specific metabolic flux, in a way that we can clearly define its lower and upper bounds (Fig 3A). Candidate targets can be identified based on the boundaries of projected Pareto surface. After that, we examined how the upper bound of metabolic flux varies with cell growth rate. If the upper bound decreases with growth rate, activation of this enzyme would lead to larger Pareto deviation for flux configurations with higher growth rate, thus conferring selectivities on cells with different growth rates (Fig 3A). Conversely, inhibition of an enzyme would impair the viability of fast-growing cancer cells, if the lower bound rises monotonously with their growth rates (Fig 3A). A correlation-based monotonousness score was defined to assess the tendency of declining upper bound or rising lower bound (Materials and Methods). The full criteria for target identification were summarized in Fig 3B. The requirement that most of the individualized models for NCI-60 cell lines locate close to the boundary is necessary to allow the metabolic perturbation to draw the flux configuration out of Pareto surface and confer significant impact on cell viability.

**Fig 3.**
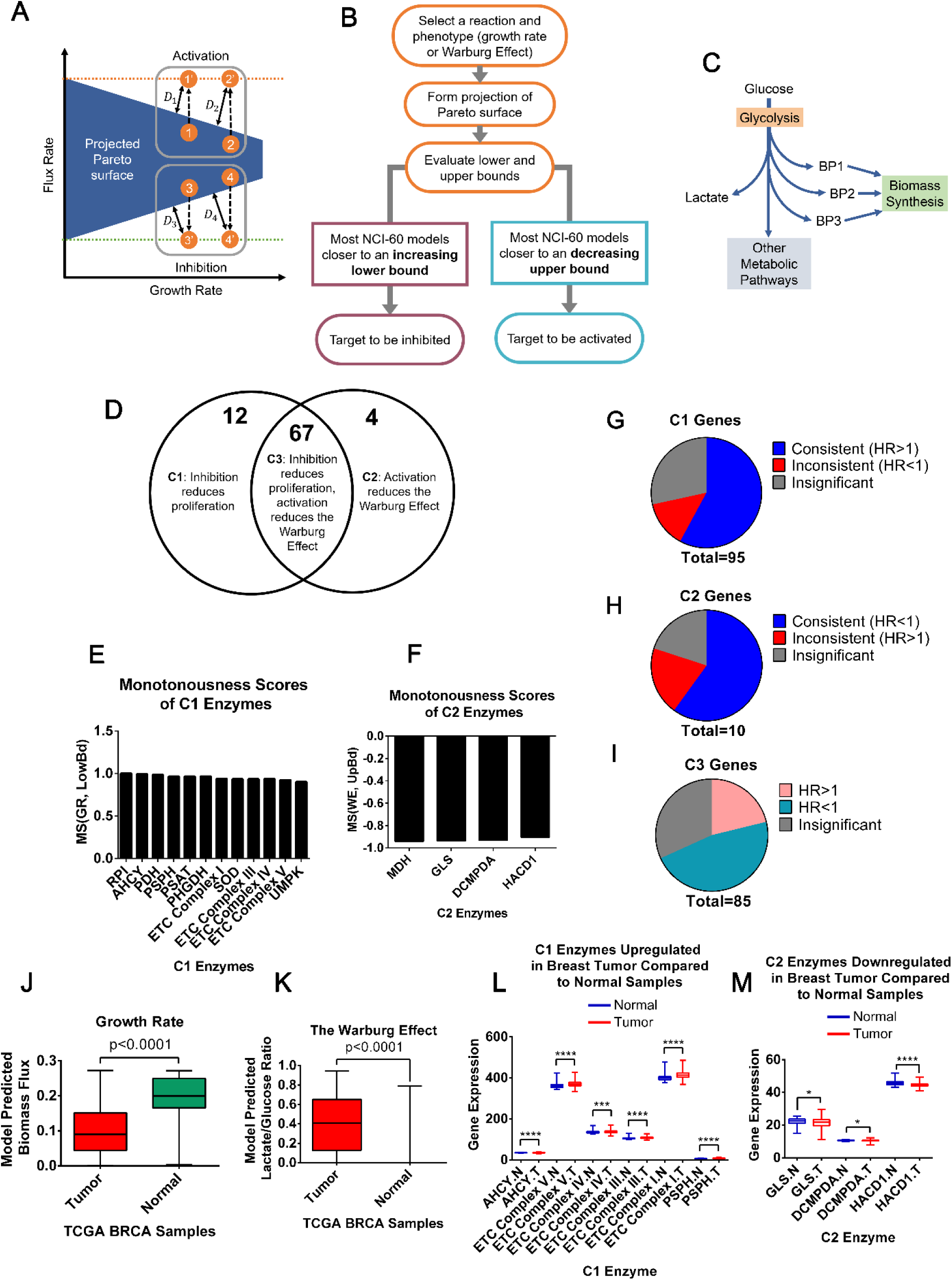
Metabolic targets identified by Pareto surface analysis exhibit strong correlations with cancer progression and patient prognosis. (A) Illustration of the criteria for target identification. (B) Workflow of identifying potential metabolic targets essential for cell proliferation and the Warburg effect. (C) A simplified illustration of glucose utilization in biomass synthesis and lactate production. (D) Summary of different categories of identified targets. (E) C1 enzymes and their monotonousness scores. (F) C2 enzymes and their monotonousness scores. (G)-(I) Results of Kaplan-Meier survival analysis of genes encoding C1, C2 and C3 enzymes. (J) and (K) Distribution of biomass production and extent of the Warburg effect predicted by the individualized models of breast tumor and adjacent normal tissues based on their gene expression profiles in TCGA. (L) Expression levels of genes associated with C1 enzymes in breast tumors and adjacent normal tissues. (M) Expression levels of genes associated with C2 enzymes in breast tumors and adjacent normal tissues. Significance levels: *: p<0.05; **: p<0.01; ***: p<0.001; ****: p<0.0001.

By analyzing the geometry of projected Pareto surface as we introduced above, we identified a list of targets essential for cancer cell growth and the Warburg Effect. The extent of the Warburg effect was quantified as the flux ratio of lactate secretion to glucose consumption. Interestingly, we found that most targets capable of reducing the Warburg effect need to be activated, whereas those able to suppress cell proliferation need to be inhibited (Table 1). Complete information about these identified targets is listed in Table S3. It has been noted that cell proliferation is a highly orchestrated process in which several pathways function in concert, producing multiple metabolic precursors for biomass synthesis (illustrated as BP1, BP2 and BP3 in Fig 3C). Defects in these pathways would prevent cells from obtaining critical building blocks, and inhibiting key enzymes involved in these processes may suppress cell proliferation. On the other hand, the Warburg effect is largely controlled by lactate dehydrogenase (LDH) generating lactate from pyruvate, which diverts carbon flux away from oxidative glucose metabolism and other related pathways. Therefore, activation of enzymes capable of consuming carbon atoms other than glycolysis may compete with the LDH flux for glucose-derived carbon atoms, leading to the reduced Warburg effect.

**Table 1.** Numbers of identified targets

|  | **Targets to be activated** | **Targets to be inhibited** |
| --- | --- | --- |
| **Inhibit cell proliferation** | 10 | 79 |
| **Inhibit the Warburg effect** | 71 | 1 |
| **Inhibit both** | 0 | 0 |

By cross-comparing the key metabolic enzymes controlling cell proliferation and the Warburg effect, we found that enzymes whose inhibition is predicted to reduce cell proliferation overlap significantly with enzymes whose activation suppresses the Warburg effect (Fig 3D). Moreover, no versatile target was predicted to exist whose perturbation is able to inhibit both processes. This result seems to be contradictory to several previous studies(Le et al., 2010; Xie et al., 2014). However, it is worth mentioning that our modeling results only reflect the direct consequence of metabolic perturbation. Metabolic enzymes often carry essential non-metabolic functions, and inhibition of cell proliferation may lead to metabolic shifts secondary to growth arrest, which were not considered by our theoretical analysis. Nevertheless, the predicted enzymes whose expression are critical for cell proliferation and/or the Warburg effect may serve as a potential target pool for therapeutic intervention, if their expression significantly correlate with disease progression.

To validate the association between identified targets and cancer progression, we conducted Kaplan-Meier survival analysis on thousands of breast cancer patients to systematically evaluate the connection between these enzymes and patient survival(Gyorffy et al., 2010). We focus on the enzymes whose activation were predicted to reduce the Warburg effect and enzymes whose inhibition would inhibit proliferation, and classify all these targets into three categories (Fig 3D):

**C1**: Inhibition of these targets was predicted to inhibit cell proliferation while no association with the Warburg effect was predicted;

**C2**: Activation of these targets was predicted to suppress the Warburg effect while no association with cell proliferation was predicted;

**C3**: Inhibition of these targets was predicted to inhibit cell proliferation, and activation of them to suppress the Warburg effect.

C1 and C2 enzymes and their corresponding monotonousness scores are shown in Fig 3E and 3F.

For C1 enzymes, we predict that their higher expression associate with worse disease progression, thus correlating with poorer prognosis and higher risk of death (*i.e.* hazard ratio>1 in Kaplan-Meier analysis). Similarly, higher expression of C2 enzymes are expected to associate with lower risk of death (hazard ratio<1 in Kaplan-Meier analysis). The correlation between C3 enzymes and patient survival may not be significant, since their roles in regulating cell proliferation and the Warburg effect counteract with each other. Kaplan-Meier survival analyses strongly support these conclusions (Fig 3G-3I).

We further conducted Kaplan-Meier analyses of C1 and C2 enzymes using gene expression and clinical datasets for non-small-cell lung cancer(Gyorffy et al., 2013) and observed similar results (Fig S5). The consistency between model-predicted functions of C1 and C2 enzymes and their association with cancer prognosis implies that these two categories of enzymes are potential anti-tumor targets. Although it is still challenging, recent progress in designing enzymatic agonists(Cool et al., 2006; Meng et al., 2016) makes C2 targets potentially druggable in developing new cancer therapeutics. On the other hand, C1 enzymes are more likely to be feasible targets, since their down-regulation is associated with better prognosis.

Regarding C3 enzymes, their overexpression may suppress the Warburg effect, whereas their lower expression may inhibit cell proliferation. Interestingly, we found that the up-regulation of most C3 enzymes correlate with better patient prognosis, indicating that their roles in regulating the Warburg effect are more important during tumor progression (Fig 3I,S5C). To further test this idea, we constructed individualized models for 1101 invasive breast carcinoma (BRCA) tumors and 114 adjacent normal tissue samples based on their gene expression profiles in the Cancer Genome Atlas(Cancer Genome Atlas, 2012), and used them to predict the proliferation potential and extent of the Warburg effect of corresponding samples. Indeed, tumor samples were predicted to exhibit significantly higher glycolysis than normal tissue samples (p=1.34×10^-41^, Wilcoxon’s rank sum test, Fig 3K), whereas the biomass production fluxes of tumor samples were predicted to be lower (Wilcoxon’s rank sum test p=1.28×10^-35^, Fig 3J). These results confirm that the Warburg effect, rather than rapid proliferation, is a more distinguished feature for malignant tumors, although targeting cell proliferation may still be considered as a therapeutic option, especially for early-staged cancers. Consistent with this notion, we observed the up-regulation of a large number of C1 enzymes (Fig 3L) and down-regulation of most C2 enzymes (Fig 3M) in tumor relative to normal tissue samples. Taken together, these integrated analyses of omics datasets validated the close association between model-predicted anti-tumor targets and cancer progression.

### Ablation of C1 enzymes impairs cancer cell proliferation

To further validate our modeling method, we focused on the top five C1 targets (Fig 3D) (Materials and Methods) and subjected four of them (RPIA, AHCY, PHGDH, and PSAT1) to experimental validation. PSPH was omitted because it catalyzes the very last reaction of serine anabolism(Locasale, 2013), thus is identical to PSAT1 as a target for our analysis. Most cell lines used in our experiments were selected from the NCI-60 panel, whereas some unavailable lines were replaced by alternatives with identical cancer types (Table S6). For each specific target, the top four cell lines with the largest growth reduction upon target inhibition as predicted by our model were selected for experimental validation, and HeLa was also included as a common cancer cell line. Efficiencies of gene ablations were validated by quantifying the mRNA levels using RT-PCR (Fig 4A-4D) and protein levels using Western blot (Fig S6). Indeed, knockdown of each individual target was associated with strong anti-proliferative effects in almost all cell lines predicted to be sensitive for target inhibition (Fig 4E-4H).

**Fig 4.**
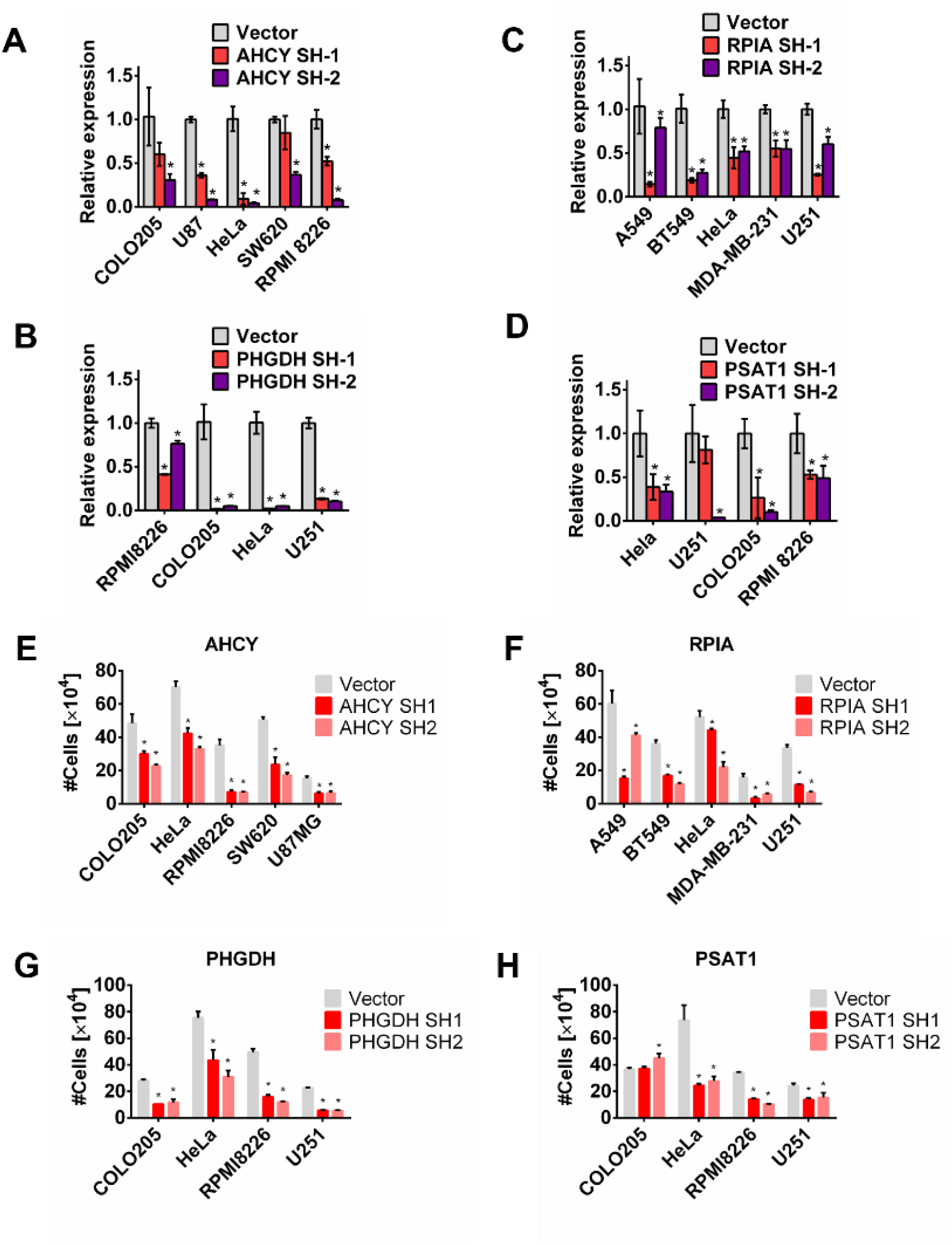
Knockdowns of C1 enzymes impair cancer cell proliferation. (A-D) Relative mRNA expression levels of AHCY, RPIA, PHGDH and PSAT1 upon shRNA knockdown in the tested cell lines. (E-H) Number of cells after 4 days upon shRNA knockdown of AHCY, RPIA, PHGDH and PSAT1 in the tested cell lines.

### Knockdown of C1 enzymes fail to inhibit the Warburg effect in cancer cells

A key prediction of our theoretical model is that goals of inhibiting cell proliferation and the Warburg effect largely conflict with each other. In other words, there is few perturbation strategies on a single enzyme capable of inhibiting cell proliferation and the Warburg effect simultaneously. To validate this conclusion, we measured lactate secretion rates with or without C1 knockdown in cell lines whose proliferation rates exhibit significant reduction upon C1 ablation. Consistent with our model, lactate production rates were not reduced in these cell lines (Fig 5A-5D).

**Fig 5.**
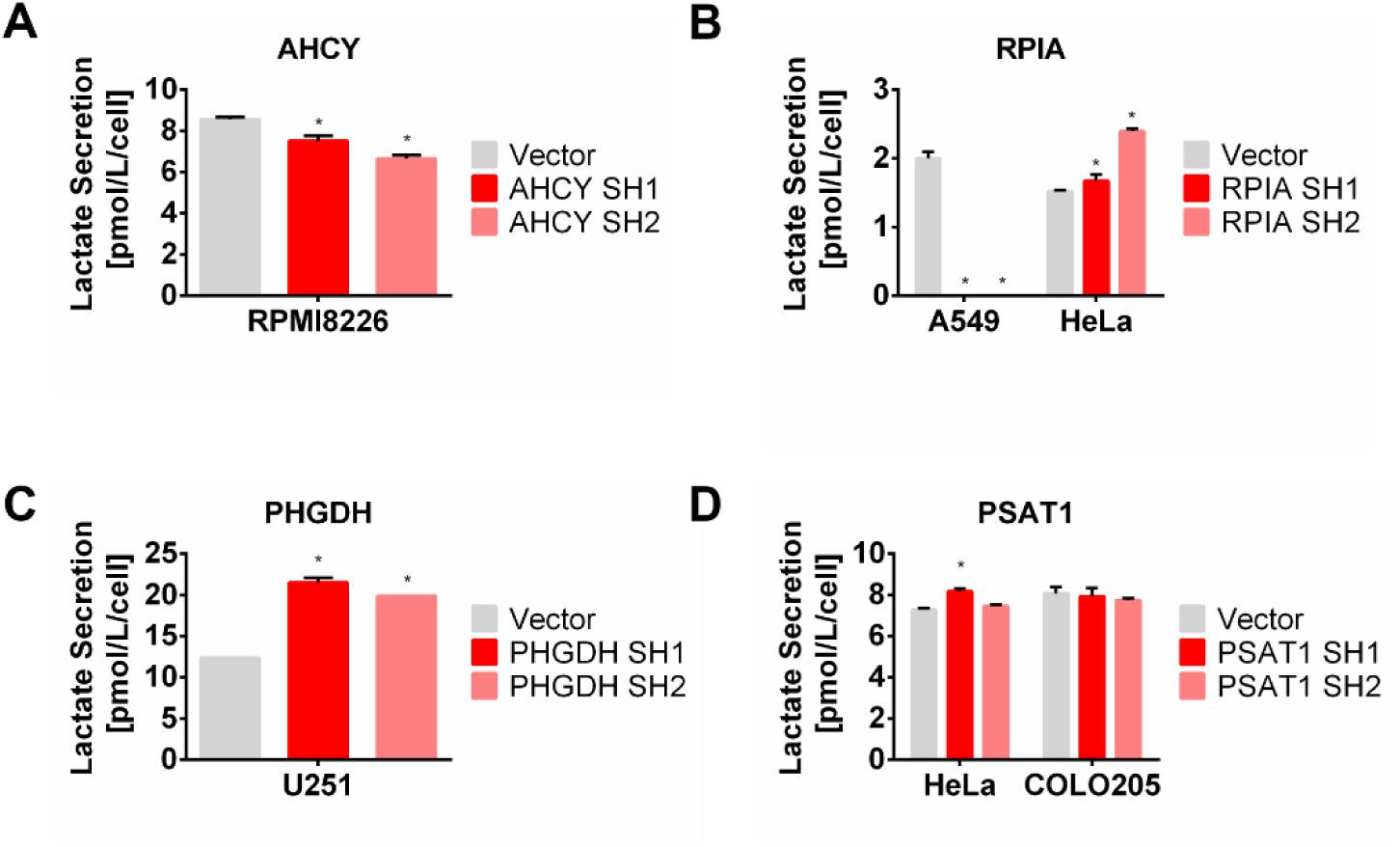
Knockdowns of C1 enzymes have minimal effect on the Warburg effect. (A) Effects of AHCY knockdown on lactate secretion. (B) Effects of RPIA knockdown on lactate secretion. (C) Effects of PHGDH knockdown on lactate secretion. (D) Effects of PSAT1 knockdown on lactate secretion.

## Discussion

### A multi-objective optimization model correctly predicts cancer cell responses to metabolic perturbation

In this study, we developed a novel strategy to model metabolism based on the assumption of multi-objective optimization. Specifically, we applied the concept of Pareto optimality to predict flux configurations with optimality in balancing the maximization of yields (growth and energy) and minimization of costs (enzymes and nutrients). By integrating these metabolic objectives with multi-omics datasets, we were able to construct individualized models to correctly predict multiple phenotypes of cancer cells including cell growth rates and responses to metabolic perturbation. This is the first attempt, to our best knowledge, to incorporate multiple objectives in modeling cancer metabolism, which demonstrates that calculated deviations from Pareto optimality with different goals closely resemble impairments of cell viability.

In our current model, we selected 4 most commonly utilized metabolic objectives for FBA analysis, including maximization of biomass production, maximization of ATP hydrolysis, minimization of total abundance of metabolic enzymes, and minimization of total carbon uptake. Nevertheless, some other objectives may also be considered for quantifying the metabolic network, such as minimization of redox imbalance, maximization of resistance to cytotoxic agents, minimization of reactive oxygen species (ROS) production, etc. Incorporating additional objectives in our model may further improve the fitting accuracy of Pareto surfaces to the actual metabolic network under different circumstances. Strategies to deduct the best combinations of objectives(Hart et al., 2015; Zhao et al., 2016) may be combined with our modeling method, and provide new insights in the reprogramming mechanism of cancer metabolism.

### The landscape of Pareto surface implicates roles of metabolic enzymes in cancer progression

Our theoretical model successfully dissects the cancer metabolic network and identifies its vulnerabilities from a global perspective. More specifically, we were able to determine several key metabolic targets to control cancer cell proliferation or the Warburg effect by analyzing the geometry of Pareto surface. Most of the potential targets associate with patient survival and exhibit differential expression patterns in cancer and normal tissues. Specifically, we identified 12 enzymes whose down-regulation exhibit strong inhibitory effects on cell proliferation yet no inhibitory impact on the Warburg effect (C1 enzymes). These enzymes are involved in multiple metabolic pathways including the Krebs cycle, oxidative phosphorylation, de novo serine synthesis, pentose phosphate pathway, and pyrimidine metabolism. Among these 12 enzymes, 11 were confirmed to correlate with cancer progression (Table 2), and some of them already have chemical inhibitors subjected to clinical tests. The clear enrichment of known anti-tumor targets in C1 enzymes, together with their significant association with patient survival, highlight the ability of Pareto optimality-based strategy in unraveling the metabolic vulnerabilities of cancer cells. Finally, we validated the top 4 C1 enzymes in human cancer cell lines with shRNA knockdown, confirming that inhibition of these enzymes would significantly reduce cell growth rates but have no inhibitory effect on aerobic glycolysis. Taken together, our theoretical and experimental results suggest that roles of metabolic enzymes in cancer progression can be uncovered by analyzing the landscape of Pareto surface under the framework of four-objective optimization model.

**Table 2.**
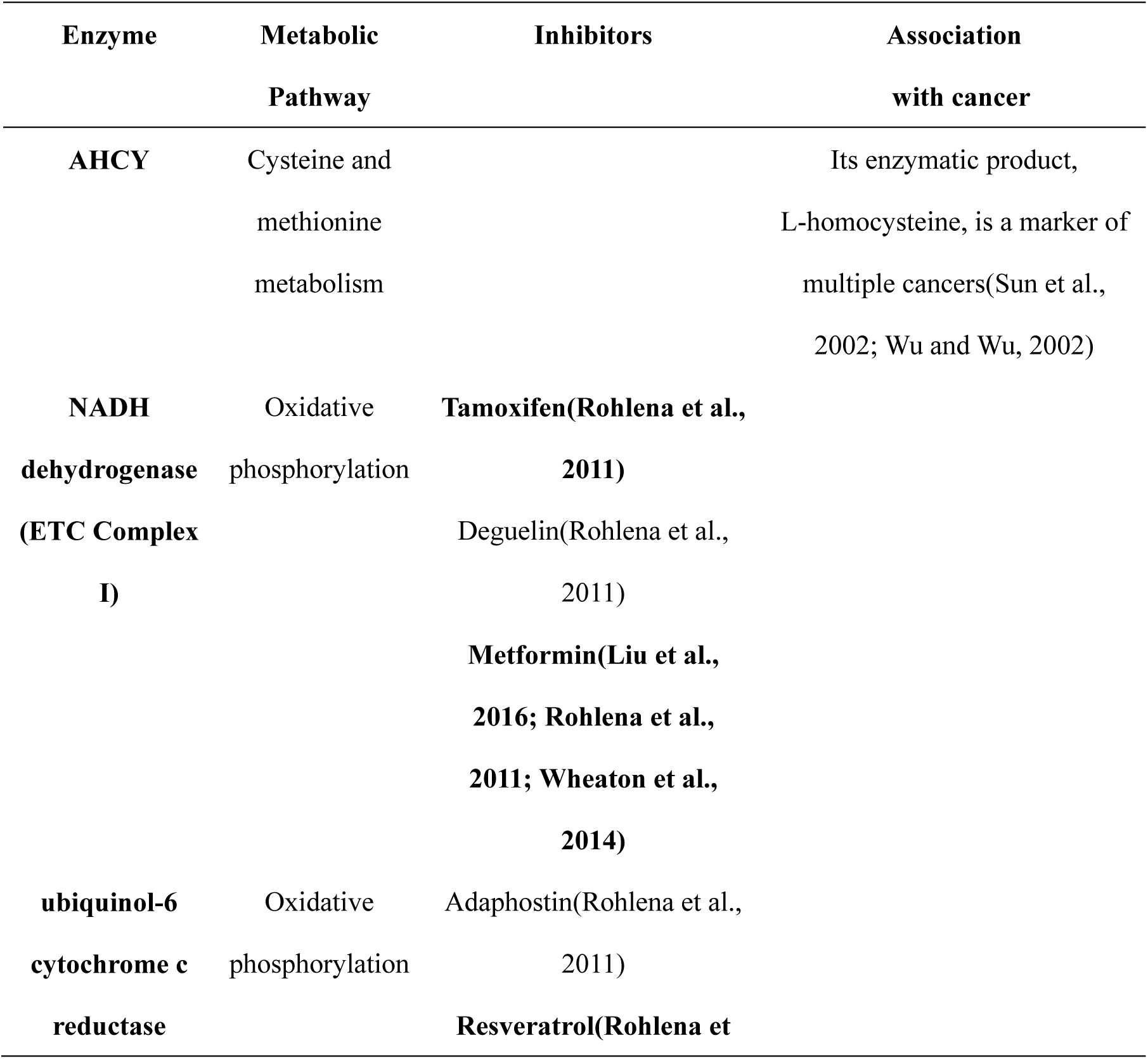
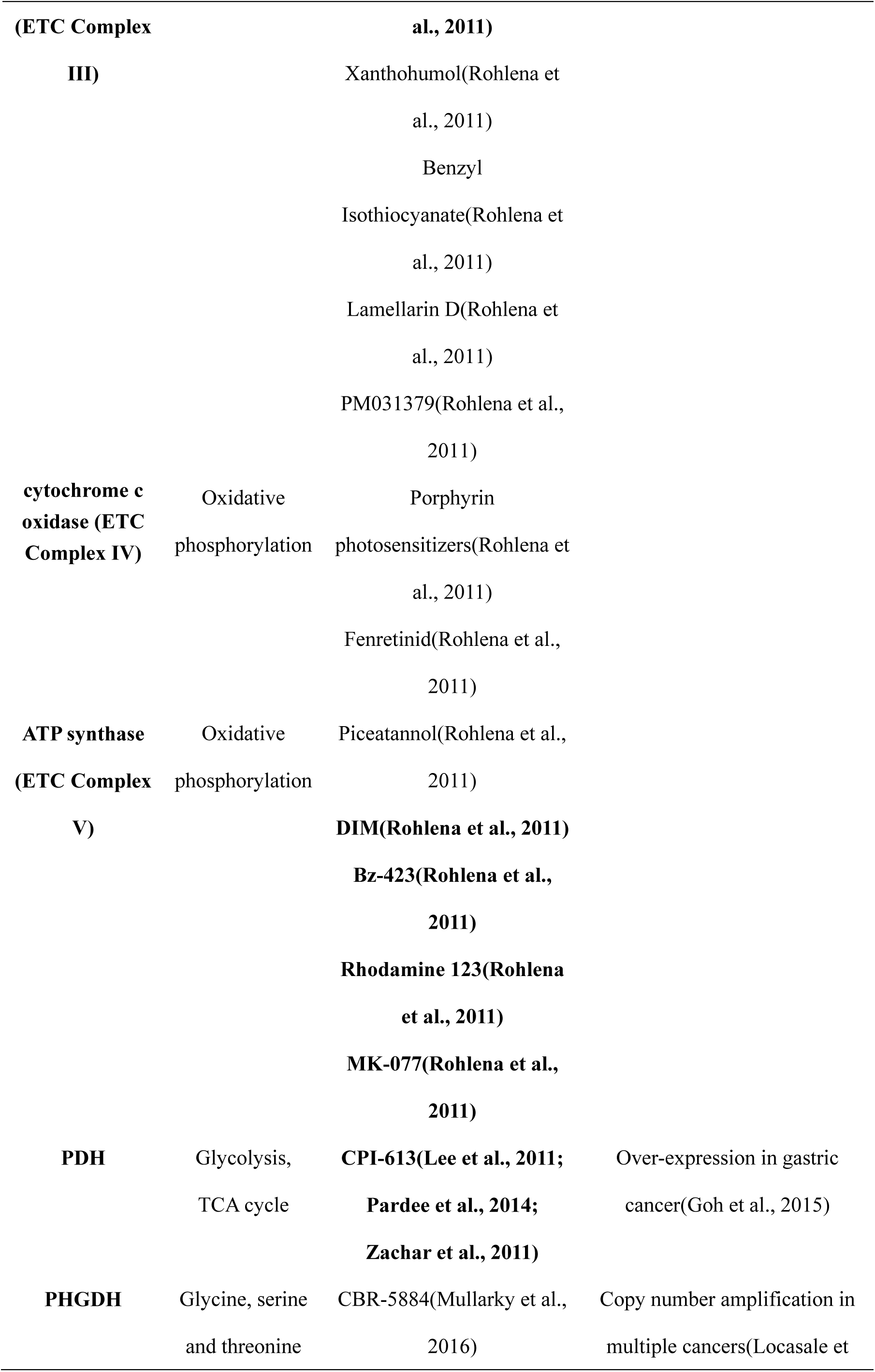
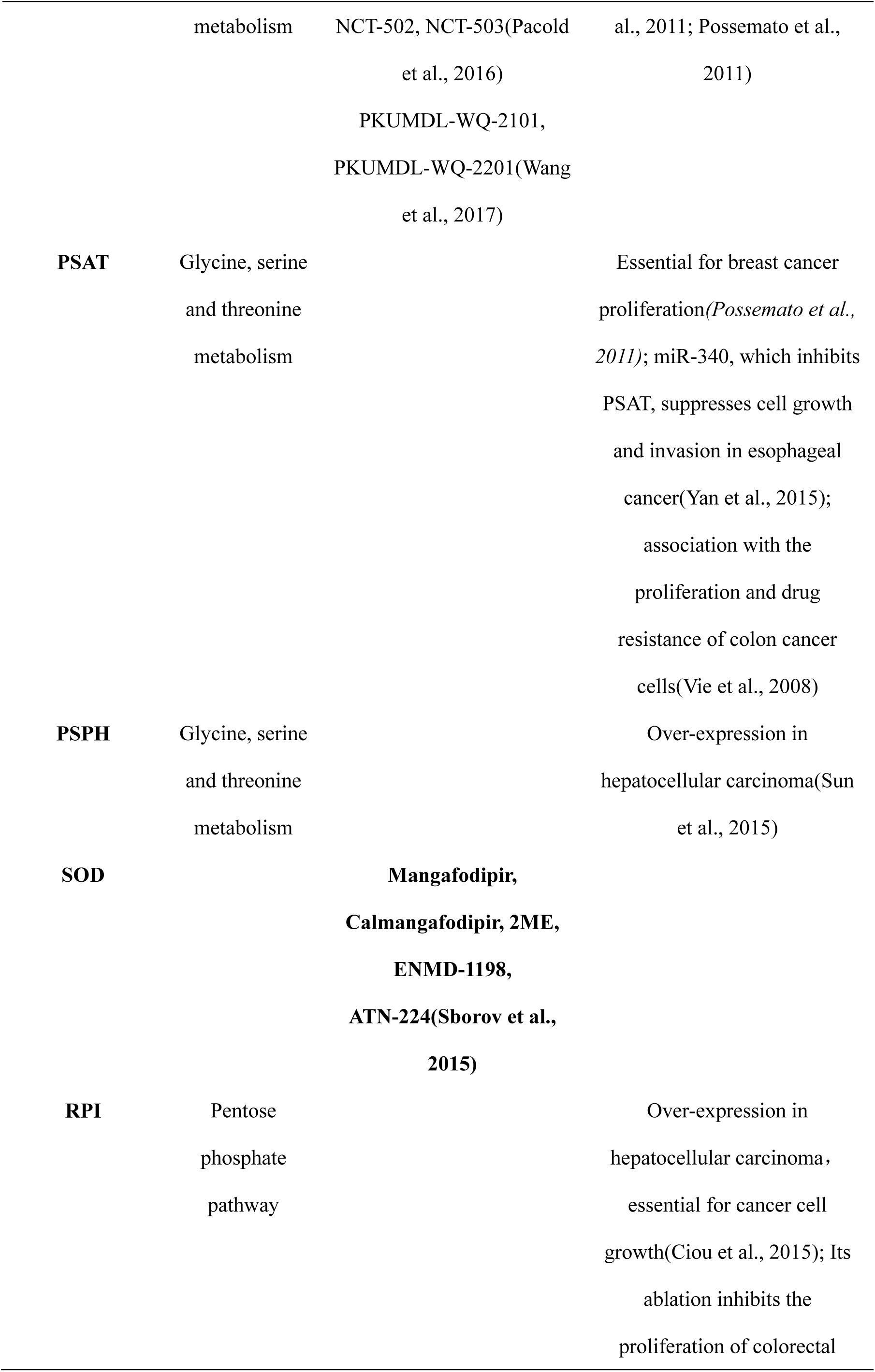
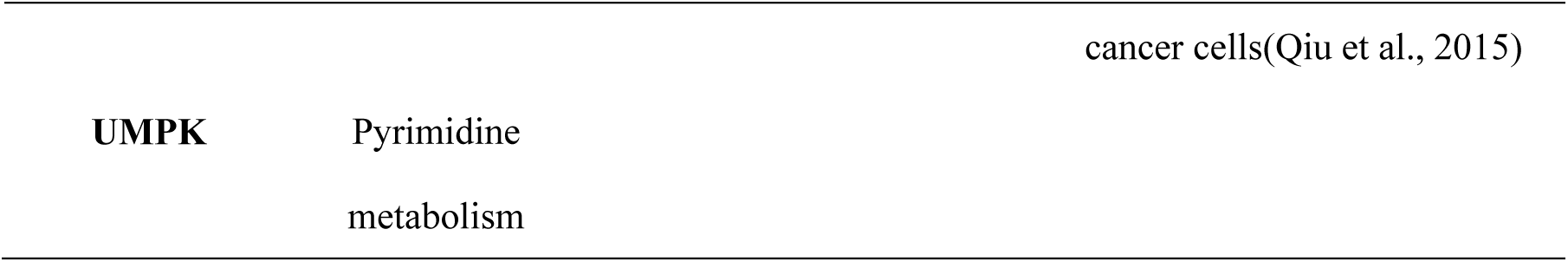
Summary of C1 enzymes

### The multi-objective optimization model sheds new light on designing anti-tumor therapeutics

Our study also suggests a novel route to target cancer-specific metabolic abnormalities by activating metabolic enzymes to compete with the Warburg effect. Although suppressing the Warburg effect has been intensely studied as a promising cancer therapy, most related strategies focus on direct or indirect inhibition of enzymes associated with aerobic glycolysis. One example is PDK1, the inhibition of which would attenuate its inhibitory effect on PDH, and facilitate oxidative glucose metabolism(Galluzzi et al., 2013). Interestingly, PDH was predicted as a C1 enzyme by our model, due to the longer distance between the individualized models and the upper bound of projected Pareto surface for the Warburg effect, as compared to the distance between the NCI-60 models and lower bound for cell growth rate, suggesting that PDH is more likely to be rate-limiting for proliferation. Consistent with this prediction, PDH inhibition has been reported to suppress cell proliferation (Table 2). Except for PDH, our analysis also reveals the wide existence of enzymes whose activation may impair tumor development mainly by inhibiting the Warburg effect. Compared to traditional methods that directly target glycolysis, this strategy may greatly reduce the side effects in normal cells that also use glucose as the major energy source. Therefore, this approach warrants further investigation even though efforts may be taken to generate enzymatic agonists.

Moreover, our model highlighted a contradictory role played by several metabolic enzymes in affecting cell growth and the Warburg effect. For a group of enzymes identified as potential targets for rapid proliferation, their activations were predicted to inhibit aerobic glycolysis (C3 enzymes in Figure 3D). The conflict between inhibiting cell proliferation and the Warburg effect reflects the intrinsic robustness of cancer as a complex disease, and was further supported by the fact that ablations of C1 enzymes failed to impair lactate production in most tested cancer cells. However, this could also be due to the fact that our modeling approach only considers the direct influence of metabolic perturbation, not the secondary effects derived from primary manipulations. In addition, our method only incorporated the stoichiometric constraints of metabolic fluxes, and ignored nonlinear factors such as the allosteric regulation of metabolic enzymes for modeling feasibilities. Further investigation is needed to characterize precise roles of those enzymes in cancer. Nevertheless, our study presented a comprehensive strategy to identify cancer-associated vulnerabilities with much-improved accuracies, as supported by survival analyses and cell-based experiments.

In summary, we have developed a novel method to model cancer metabolism based on Pareto optimality under the framework of multi-objective optimization. This approach created an integrated workflow from omics-based mathematical model to metabolic target identification, and predicted metabolic hubs essential for cancer cell proliferation and/or the Warburg effect. The high consistency between predicted roles of metabolic enzymes in cancer and tumor ‘omics’ data suggests that the overall effect of a specific enzyme during tumor development is determined by its functions in multiple metabolic tasks rather than a single task such as cell proliferation. In addition to modeling cancer metabolism, this methodology may also be applied to explore other disease-related metabolic abnormalities with accessible omics datasets.

## Materials and Methods

### Defining metabolic objectives in the genome-scale metabolic model

We considered four metabolic objectives including maximization of biomass production flux (f_BM_), maximization of ATP turnover (f_ATP_), minimization of carbon uptake (CU), and minimization of total metabolic enzyme abundance (EA). The genome-scale metabolic model used here, Recon 1, already contains a biomass producing flux whose coefficients are determined by the molecular composition of mammalian cells. We employed a curated and decomposed model by Shlomi *et al* (i.e. all reversible reactions are decomposed into forward and backward reactions) to simplify the following procedures, but this model lacks some critical fluxes and biomass components. Therefore, we downloaded model files in the SBML format from the BioModels Database, translated them into MATLAB files using COBRA, and supplemented the original model with ignored components for the following analysis. The MATLAB file of updated model bas been included in the Supplementary Materials (Data S1). This model also contains an ATP hydrolysis flux, whose maximization was considered as the second objective. Eventually the carbon uptake (CU) flux was calculated as follows:

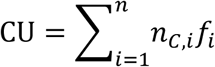

n is number of fluxes in the model, *n*_*C,i*_ is the number of carbon atoms imported into intracellular compartments by the ith flux and *f*_*i*_ is its flux rate. If this flux does not lead to any carbon uptake, the value of *n*_*C*,*i*_ is zero.

The enzyme abundance (EA) was determined by:

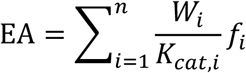

Since the coefficients 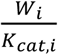 have already been evaluated in the curated model with a solvent capacity constraint, they were simply utilized in our analysis. In other words, the solvent capacity constraint has been replaced by the objective of minimizing total enzyme abundance. Upper limits of nutrient influxes were set according to the composition of RPMI-1640 medium (Table S2).

**Sampling the Pareto surface with the Branched ε-Constraint Method (BECM)**

Pareto solutions of optimization problems with N objectives can be calculated by ε-Constraint Method, in which N-1 of the objectives are transformed into soft constraints and the left single objective can be optimized with the resultant constraints. Assume that we are interested in solving the following problem:

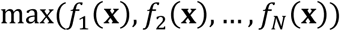

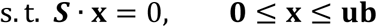

In which **S** is the stoichiometric matrix, **x** is the flux configuration in which each element is flux rate of a reaction, **ub** is the vector of maximal rates of reactions. Assume that *f*_1_(**x**) is the objective to be optimized and all other objectives are treated as constraints, the derived single objective optimization problem (SOOP) becomes:

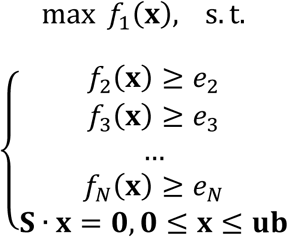

By transforming different combinations of objectives to constraints, adjusting the constants e_1_ to e_n_, and repeatedly solving the corresponding optimization problem, we can obtain a solution set with Pareto optimality. In our model all objectives and constraints are linear, enabling us to solve the SOOPs with efficient algorithms (e.g. simplex method, interior point method, etc) for linear programming (LP). The LP problems were solved using Mosek (http://www.mosek.com/).

To generate feasible SOOPs more efficiently, we employed a branched strategy. First we calculate the maximal values for the biomass production flux 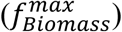 and the ATP hydrolysis flux 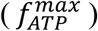. We then sampled 10000 combinations of (*e*_*Biomass*_, *e*_*ATP*_) in the region 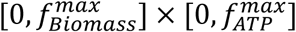 with Latin Hypercube Sampling. For each combination we calculated the minimal values of CU and EA (*CU*_*min*_ and *EA*_*min*_) compatible with constraints in the GSMM, *f*_*Biomass*_ ≥ *e*_*Biomass*_ and *f*_*ATP*_ ≥ *e*_*ATP*_. Let *CU*_max_ and *EA*_max_ denote the value of CU when EA reaches its lowest limit, and the value of EA when CU reaches its lowest limit. Calculating the range of [*CU*_*min*_, *CU*_*max*_] and [*EA*_*min*_, *EA*_*max*_] according to sampled (*e*_*Biomass*_, *e*_*ATP*_) prior to SOOP generation helps to avoid generating a large number of infeasible SOOPs. We finally generated SOOPs in a branched manner in which stratified sampling were applied to select m_1_ values within [*CU*_*min*_, *CU*_*max*_] and m_2_ values within [*EA*_*min*_, *EA*_*max*_]. m_1_ + m_2_ SOOPs were generated according to m_1_ minimizing EA and m_2_ minimizing CU. The values of m_1_ and m_2_ were determined by the range of [*CU*_*min*_, *CU*_*max*_] and [*CU*_*min*_, *CU*_*max*_] with 10 as the maximal value. Possible solutions of all feasible SOOPs (42930 in total) were summed up as the sampled Pareto surface.

### Retrieving and processing the omics datasets

Proteomics data of the NCI-60 cell lines are available at the NCI-60 Proteome Resource: http://wzw.tum.de/proteomics/nci60. The exchange flux rates of these cell lines were presented in a study at 2012(Jain et al., 2012). However, these proteomics and fluxomics data were only normalized by cell numbers, and NCI-60 cell lines were noted to exhibit different cell sizes. Therefore, we further normalized the original data by cell volumes calculated based on cell diameters available at: http://www.nexcelom.com/Applications/Cancer-Cells.html, with the assumption that single cells are perfect spheres. Expression levels of metabolic enzymes were evaluated by gene-protein-reaction rules included in the GSMM (Details in Supplementary Materials). Gene scores quantifying sensitivities to gene knockdown are available at: http://www.broadinstitute.org/achilles. Cell growth rates are available at: http://discover.nci.nih.gov/cellminer/. The TCGA datasets for breast cancer are available at the Cancer Genome Browser (https://genome-cancer.ucsc.edu)(Zhu et al., 2009). Kaplan-Meier survival analyses were performed using the Kaplan-Meier Plotter (http://kmplot.com/analysis/)(Szasz et al., 2016).

### Constructing the individualized models

We defined a similarity score to evaluate to which extent the distribution of flux configurations as predicted by the individualized models can reproduce the distribution of corresponding omics data. The similarity metric (S) is comprised of two parts: *S*_1_ and *S*_2_. *S*_1_ is the summation of correlation coefficients between model-predicted and experimentally-determined exchange flux rates:

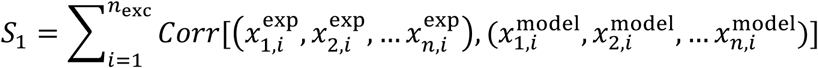

*S*_2_ is the summation of correlation coefficients between model-predicted average flux rates and average expression levels of metabolic pathways:

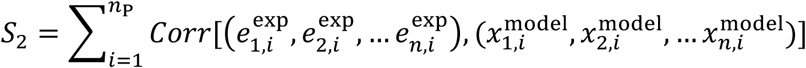

*n*_exc_ is the number of exchange fluxes whose measurements were available, *n*_P_ is the number of metabolic pathways in the KEGG Database, n is the number of NCI-60 cell lines whose fluxomics and proteomics data are both available, 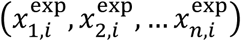 is the experimentally-determined flux rates of cell lines for the ith exchange flux, 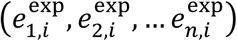 is the average expression levels of metabolic enzymes for the ith pathway according to proteomics data, 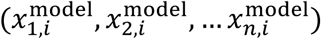 is the flux rates predicted by our models (individual fluxes in *S*_1_ and pathway-averaged fluxes in *S*_2_). The correlation metric used here is Spearman’s rank correlation coefficients. Single enzymes were mapped to metabolic pathways according to the BRENDA Database based on their EC Numbers. The individualized models (i.e. flux configurations for the cell lines) were constructed along the approximate Pareto surface by maximizing the similarity metric S with simulated annealing. For the TCGA datasets in which no flux rates are available, the similarity score S_2_ was maximized alone.

**Simulation of metabolic gene ablations and calculation of the Pareto deviation score**

We simulated the effects of metabolic gene knockdowns with minimization of metabolic adjustments (MOMA)(Segre et al., 2002). First we assumed that all metabolic genes have equal expression levels of 1 and evaluate the expression levels of all enzymes contained in the GSMM. We then changed the expression level of to-be-ablated gene to zero and re-evaluated this enzyme. For all reactions associated with enzymes whose expression levels change in this process, let *E*_0_ and *E*_1_ denote the evaluated expression levels of the corresponding enzyme before and after the knockdown, respectively. After that, we adjusted the upper bound of flux through this reaction by multiplying itself with *E*_1_/*E*_0_. Finally, the new flux configuration **x**_1_ after gene knockdown was calculated by minimizing the Euclidean distance to the original flux configuration **x**_0_ with the new upper bound constraints. Pareto deviation score (PDS) was then computed as below:

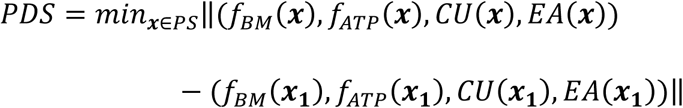

### Analyzing the projected Pareto surface

Briefly, the upper and lower bounds of projected Pareto surface were approximated by mathematical discretization. Ranges of cell growth rate or the Warburg effect (defined as the ratio of lactate secretion flux to glucose uptake flux) of all Pareto solutions were divided into 100 bins with identical size. Let *f*_*i*_ denote the variable describing the ith flux used in combination with cell growth rate or the Warburg effect for the projection, the lower and upper bounds of *f*_*i*_ in all Pareto solutions whose growth rates or the Warburg effect fell in each of the 100 bins were calculated as [*LB*_1_, *LB*_2_,…, *LB*_100_] and [*UB*_1_, *UB*_2_,…, *UB*_100_]. The tendency of these bounds to be monotonous (increasing or decreasing) were quantified by Spearman’s rank correlation coefficients between the vector [*LB*_1_, *LB*_2_,…, *LB*_100_] or [*UB*_1_, *UB*_2_,…, *UB*_100_] and the vector [1, 2, 3, …, 100]. We refer to these correlation coefficients as monotonousness scores (Table S5). Bounds were considered as monotonously decreasing if correlation coefficients were less than −0.9, and monotonously increasing if larger than 0.9. If a projection has decreasing upper bound and more than half of the individualized models for NCI-60 cell lines are located closer to the upper bound than to the lower bound, the enzyme catalyzing the flux used for projection would be identified as a potential target to be activated. Conversely, increasing lower bound to which more than half of the individualized models are located closer would identify a potential target to be inhibited.

### Experimental validation of identified metabolic targets

Cell lines were selected from the NCI-60 panel based on their predicted changes of biomass production flux using the NCI-60 individualized models as previously constructed. The simulation of enzymatic perturbations was performed by MOMA. Cell lines predicted to have significant reduction of biomass production flux were selected for further experimental validation.

**Cell Culture.** BT549, MDA_MB_231, A549, U87, SW_620, COLO205, and RPMI_8226 cell lines were purchased from the China Infrastructure of Cell Line Resources and cultured in RPMI containing 10% FBS and antibiotics. Purchased U251 and HeLa cells were cultured in DMEM containing 10% FBS. All cell lines were confirmed to be mycoplasma negative. shRNA constructs were transfected into cells using Lipofectamine and selected with corresponding antibiotics.

**Immunoblot Analysis.** Cells were lysed with lysis buffer (25 mM Tris, 100 mM NaCl, 1% Triton X-100, 1 mM EDTA, 1 mM DTT, 1 mM NaVO_4_, 1 mMb-glycerol phosphate, and 1 mg/mL aprotinin), and then the lysates were resolved by SDS-PAGE and proteins transferred to PDVF membranes. The filters were incubated with various primary antibodies diluted in TBST (20 mM Tris, 135 mM NaCl, and 0.02% Tween 20). The primary antibodies were detected with horseradish peroxidase-conjugated secondary antibodies followed by exposure to ECL reagent.

**Cell growth and metabolic assays**. Cells were plated in dishes at a density of 5×10^4^ cells/dish and cultured in low serum medium for 5 consecutive days. Every other day one set of cells was collected and counted, while the medium on the remaining sets of cells was replenished. The rates of lactate production were determined using a BioProfile basic biochemistry analyzer (Nova Biomedical).

**Statistical analysis.** Statistical tests were performed using MATLAB. Algorithms for sampling the Pareto surface, constructing individualized models, and identifying targets were implemented in MATLAB codes.

## Acknowledgement

This study was supported in part by the Ministry of Science and Technology (2016YFA0502303 and 2015CB910300 to LL), the National Natural Science Foundation of China (21633001 to LL, 81572508 to BL) and the Fundamental Research Funds for the Sun Yat-sen University (16ykzd04 to BL). ZD thanks Dr. Ning Yin for helpful discussions and insightful comments on the manuscript and Dr. Chunmei Li for help with the cancer cell lines.

